# Systematic evaluation and validation of reference and library selection methods for deconvolution of cord blood DNA methylation data

**DOI:** 10.1101/570457

**Authors:** Kristina Gervin, Lucas A. Salas, Kelly M. Bakulski, Menno C. van Zelm, Devin C. Koestler, John K. Wiencke, Liesbeth Duijts, Henriëtte A. Moll, Karl T. Kelsey, Michael S. Kobor, Robert Lyle, Brock C. Christensen, Janine Felix, Meaghan J. Jones

**Author notes:** Contributed equally to this work (Kristina Gervin, Lucas A. Salas).

## Abstract

**Background:** Umbilical cord blood (UCB) is commonly used in epigenome-wide association studies of prenatal exposures. Accounting for cell type composition is critical in such studies as it reduces confounding due to the cell specificity of DNA methylation (DNAm). In the absence of cell sorting information, statistical methods can be applied to deconvolve heterogeneous cell mixtures. Among these methods, reference-based approaches leverage age appropriate cell-specific DNA-methylation profiles to estimate cellular composition. In UCB, four reference datasets comprising DNAm signatures profiled in purified cell populations have been published using the Illumina 450K and 850K EPIC arrays. These datasets are biologically and technically different, and currently there is no consensus on how to best apply them. Here, we systematically evaluate and compare these datasets and provide recommendations for reference-based UCB deconvolution.

**Results:** We first evaluated the four reference datasets to ascertain both the purity of the samples and the potential cell cross-contamination. We filtered samples and combined datasets to obtain a joint UCB reference. We selected deconvolution libraries using two different approaches: automatic selection using the top differentially methylated probes from the function *pickCompProbes* in minfi and a standardized library selected using the IDOL (Identifying Optimal Libraries) iterative algorithm. We compared the performance of each reference separately and in combination, using the two approaches for reference library selection, and validated the results in an independent cohort (Generation R Study, n=191) with matched FACS measured cell counts. Strict filtering and combination of the references significantly improved the accuracy and efficiency of cell type estimates. Ultimately, the IDOL library outperformed the library from the automatic selection method implemented in *pickCompProbes*.

**Conclusion:** These results have important implications for epigenetic studies in UCB as implementing this method will optimally reduce confounding due to cellular heterogeneity. This work provides guidelines for future reference-based UCB deconvolution and establishes a framework for combining reference datasets in other tissues.

## Background

DNA methylation (DNAm) is involved in the regulation of genes and is essential for normal development. Human epigenome-wide association studies (EWAS) are widely used to investigate the association between DNAm variation and prenatal environmental factors, and understanding the developmental origin of phenotypes and diseases[1-3]. These studies are often performed in umbilical cord blood (UCB), which is collected from the umbilical cord after birth. UCB is easily accessible and an ideal time point for capturing and studying the influence of fetal environmental exposures on DNAm. In addition, DNAm differences in UCB can reflect systemic exposures and, in some instances, potentially serve as surrogate and proxy for tissues that cannot be easily assessed (e.g. brain tissue)[4].

Cell types present in UCB reflect those in the peripheral whole blood at birth, including hematopoietic stem cells and nucleated red blood cells (nRBCs), which rapidly decline in the newborn after birth. The median proportion of nRBCs present at birth usually ranges from 4 to 9% and rarely exceeds 22%[5]. Leukocytes in the newborn are immunologically immature, consistent with the need for these cells to develop both the appropriate response to pathogens as well as immunological memory. This is a hallmark of the adaptive immune system and is under epigenetic control[6, 7]. The major leukocyte subsets in UCB are granulocytes, monocytes, and lymphocytes, with the latter containing T cells, B cells and NK cells[8]. The different leukocytes in UCB are functionally and developmentally distinct and display cell type specific DNAm patterns[9]. In addition, different chronic and acute stressors can alter the composition of cell types in UCB between individuals. As a consequence, it is important to pay particular attention to this confounding source of variability when conducting an EWAS using heterogeneous cell mixtures such as UCB. In addition, cell type proportions can be a mediator between exposure and disease. Many EWAS have demonstrated the importance of adjusting for cell type heterogeneity[10, 11].

There are reference-free and reference-based strategies to address the problem of cell type heterogeneity in blood samples, which have been discussed and reviewed elsewhere[12-14]. A widely used deconvolution method is the reference-based using constrained projection/quadratic programming (CP/QP) proposed by Houseman et al [15]. Briefly, the CP/QP method uses a reference dataset consisting of cell type specific DNAm signatures as the basis for inferring cell proportions in samples comprised of heterogeneous mixtures of those cell types. These deconvolution estimates can then be included as covariates in the downstream statistical models to adjust for the potential confounding effects of cell type differences between samples. The CP/QP method depends entirely on a reference library consisting of cell type specific DNAm markers. Hence, a critical first step involves selection and assembly of a library that reflects a DNAm fingerprint of the cell types using a reference dataset. This is commonly performed using one of two algorithms: pickCompProbes implemented in the minfi package[16] and IDOL. In UCB, pickCompProbes [16] performs a default automatic selection process choosing the top 100 most differentially methylated probes with a F-test p<10E-08 per cell type. Compared to adult peripheral blood, applying this algorithm to UCB selects probes agnostic of the direction of DNAm difference. This leads to libraries that poorly discriminate certain leukocyte subpopulations, particularly those that have a shared lineage[17]. IDOL[18] is an iterative algorithm, which dynamically scans a candidate set of cell type specific DNAm markers for a library that is optimized to accurately estimate cell types, often referred to as leukocyte-differentially methylated regions (L-DMRs). IDOL requires a set of samples with known values for the cell mixtures, ideally artificially spiked samples with pure cell subtypes of known mixing proportions, but mixed samples with cell counts can be substituted[18, 19].

Currently, four analogous UCB references have been published consisting of cell type specific DNAm data assayed using the Illumina 450K and 850K EPIC technology[17, 20-22]. These datasets possess a range of biological and technical differences related to the number of samples, isolated cell fractions and phenotyping, purity estimates, separation method, ethnicity, sex ratio, gestational age and array technology. Although the application of one of the reference datasets for deconvolution has been validated[21], it is not known how these differences influence the deconvolution estimates. Importantly, there is no consensus on how to best use UCB reference methylome datasets, whether it is appropriate to combine reference datasets, and if so, which library selection method will have optimal performance. Therefore, the aims of this study were to: 1) give a comparative descriptive overview of the different references datasets 2) compare how different methods for selecting reference libraries impact deconvolution estimates, 3) benchmark and validate deconvolution estimates in an independent cohort containing matched cell counts and 4) provide guidelines for reference-based cord blood deconvolution using the four UCB references separately and in combinations.

## Results

### Descriptive overview and data cleaning

Four publicly available UCB reference datasets were used, named by the first author of the respective publications: *Bakulski reference, de Goede reference, Gervin reference* and *Lin reference*[17, 20-22]. These datasets possess several noticeable biological and technical differences (**Table 1**). Stringent quality control and probe filtering procedures were applied to minimize technical variation. As expected, a principal component analysis (PCA) revealed distinct clustering of the cell types according to the hematopoietic lineage (i.e. lymphoid and myeloid cells), reflecting different DNAm profiles (**Figure 1**). The references containing nRBCs (*de Goede* and *Bakulski*) demonstrate clear separation between white blood cells and erythrocytes. The CD4T and CD8T cells show very similar DNAm profiles and form partly overlapping clusters. Of note, the *Bakulski reference* also shows considerable variation not related to cell types and more overlapping cell type clusters compared to the others.

**Table 1.**
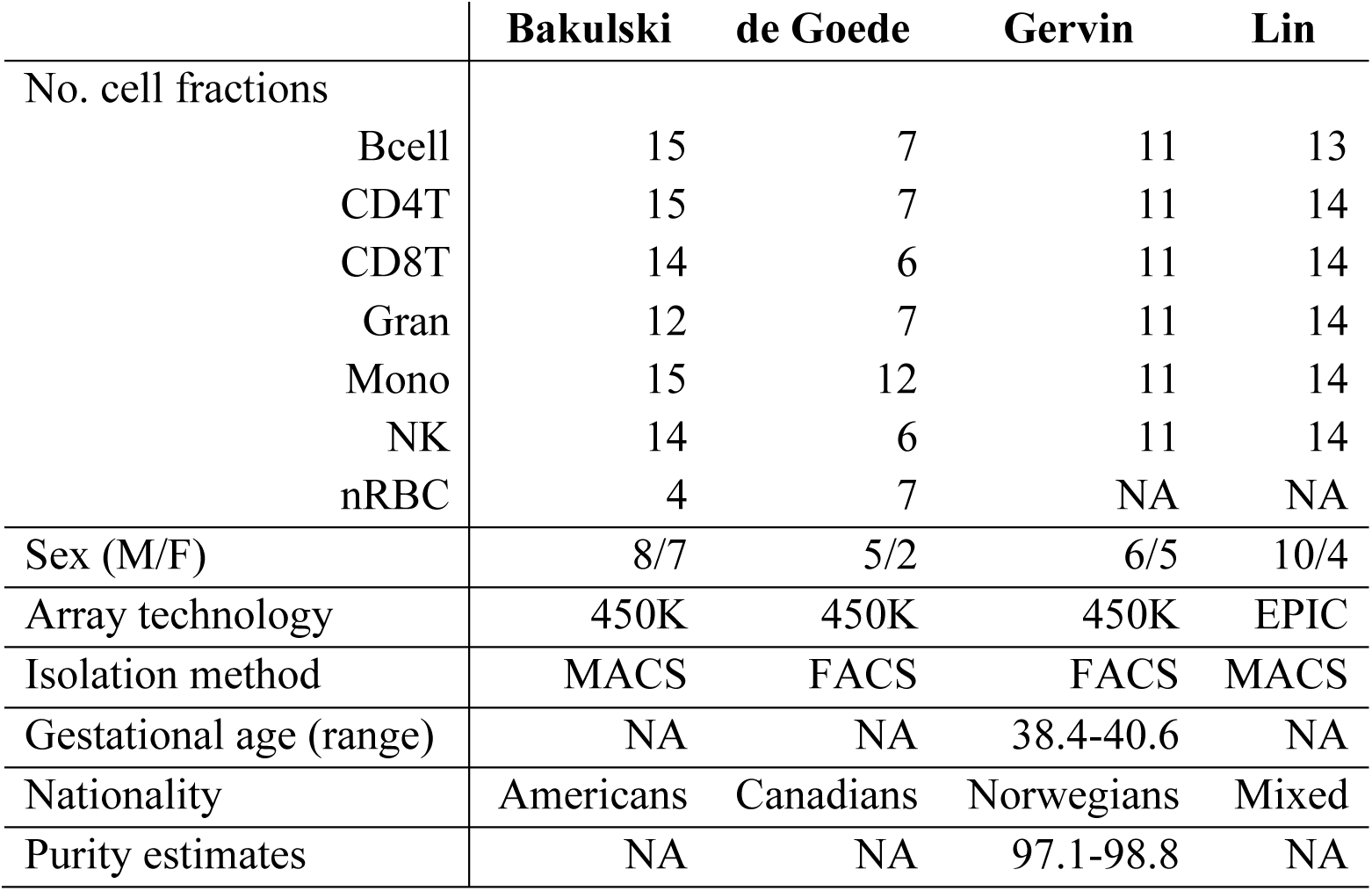
Descriptive overview of the UCB reference datasets.

**Figure 1.**
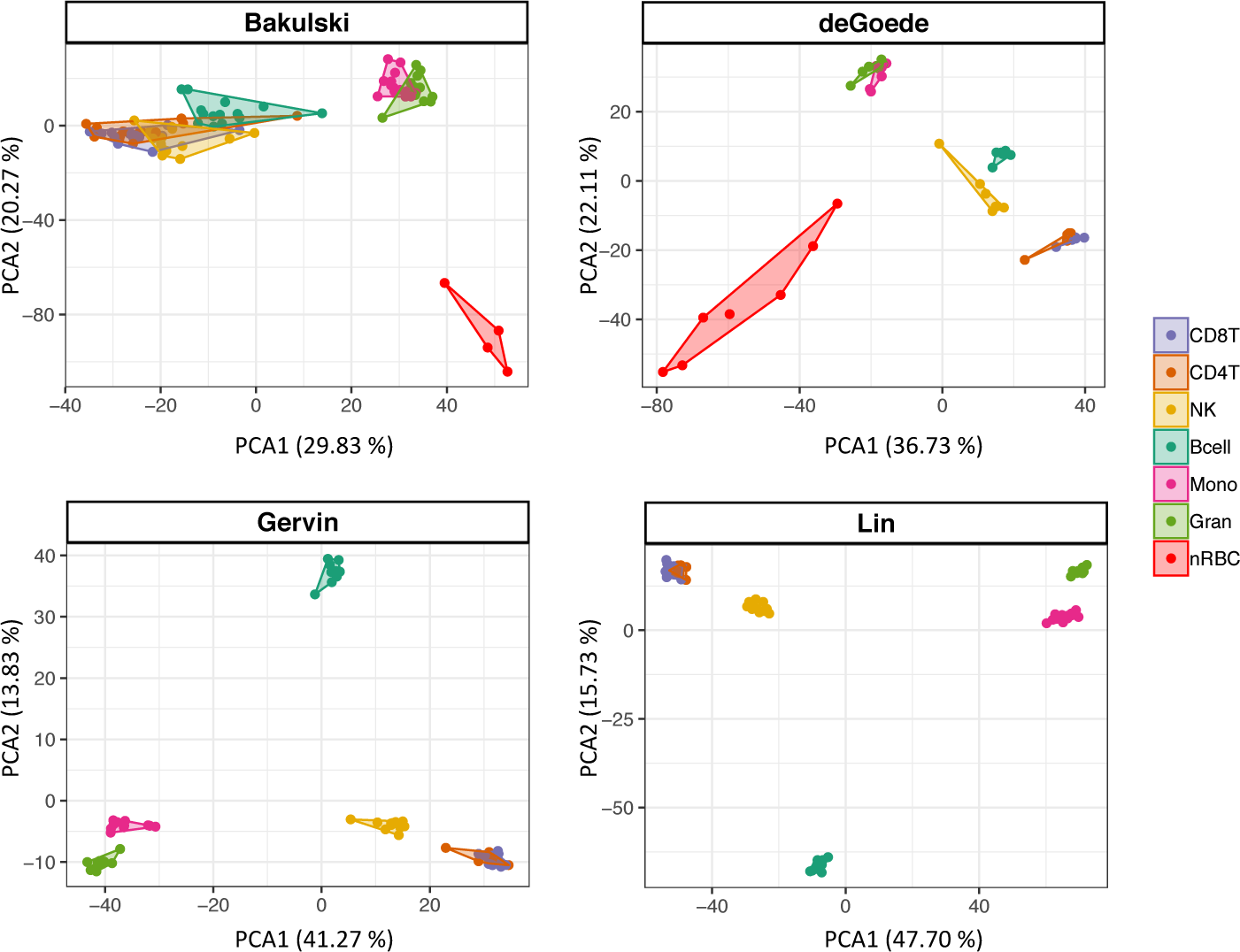
PCA scatterplot of cell type specific DNAm in four UCB references as published (raw). The two first principal components are plotted with the proportion of variance explained by each component indicated next to the axis labels. The plot clearly shows distinct clustering of the different cell types and most of the variance in DNAm can be attributed to the different cell types. Of note, nRBCs are not included in the Gervin and Lin references.

The technical and biological differences associated with each reference dataset, including purity and phenotyping (i.e. antibodies used for Magnetic-Activated Cell Sorting (MACS) and Fluorescence-Activated Cell Sorting (FACS) isolation) likely influence the deconvolution process using datasets individually, and especially in combination. To test this, we reasoned that a strict cleaning of the datasets prior to deconvolution could improve the accuracy and precision of the deconvolution estimates.

The cleaning process is described in detail in Materials and Methods. Briefly, using a projection of adult cell types from an adult reference[19], samples showing <70% of the adult cell type were removed from the reference and samples with >70% of a different cell type were reclassified to the “correct” cell type. The results from this cleaning process are shown in **Figure 2** and **Supplemental Table S1**. Using a 70% cut off resulted in removal of 24 of 89 samples (26.9%) (n=4 Bcell, n=6 CD4T, n=12 CD8T, n=1 Gran, n=1 Mono) and reclassification of 1 sample (CD8T to NK) in the *Bakulski reference.* Of note, almost all CD8T fractions in this reference showed a large proportion of NK cells (**Figure 2**). Further, 3 samples (n=3 Mono) were removed and 3 samples from the same individual reclassified to opposite cell types (n=3 CD4T to CD8T and n=3 CD8T to CD4T) in the *Gervin reference*. No samples were removed or reclassified from the *de Goede* or *Lin references*. An adult reference was not applicable for the nRBC cell type and all nRBC samples were retained. In total, 263 cell type DNAm signatures (n=42 Bcell, n=41 CD4T, n=33 CD8T, n=43 Gran, n=48 Mono, n=45 NK and n=11 nRBCs) were included in a combined cleaned UCB reference, which we have made available as an Bioconductor package named “*FlowSorted.CordBloodCombined.450K*”.

**Figure 2.**
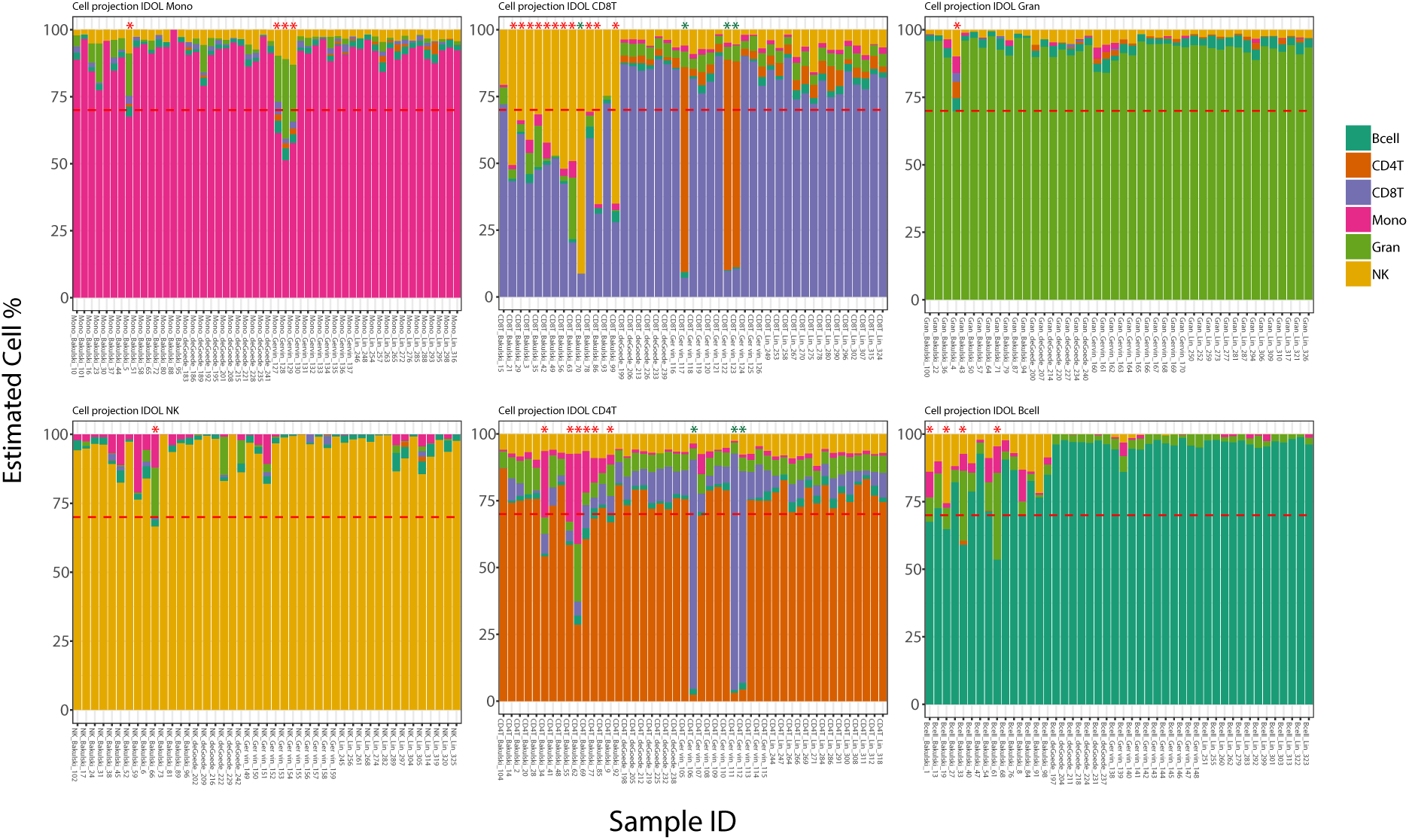
Data cleaning using a projection of adult cell types. Samples in the four UCB references showing <70% of the adult cell type were removed, whereas samples with >70% of a different cell type were reclassified to the corresponding cell type. Using a 70% cutoff resulted in removal of 24 samples (26.9%, indicated by red asterisk) and reclassification of 3 samples (indicated by green asterisk. Of note, the majority of the CD8T cell fractions in the Bakulski reference showed a large proportion of NK cells.

### Selection and comparison of reference libraries for UCB deconvolution

Deconvolution of heterogeneous cell mixtures like UCB relies on a suitable set of cell type specific DNAm markers referred to as a L-DMR library. To compare the four reference datasets (individually and in different combinations) and to evaluate how the cleaning process influenced deconvolution estimates, we used two different methods to select L-DMR libraries. These analyses were performed in a test dataset (n=24) consisting Gran, Mono, Bcell, CD4T, CD8T, NK and nRBCs UCB DNAm signatures and matched CBC and FACS counts. First, we applied the selection strategy used by pickCompProbes, which selects the top 100 differentially methylated probes per cell type (in UCB the algorithm will not select the top 50 hyper- and the top 50 hypomethylated CpGs as in adult peripheral blood). This resulted in the selection of a total number of probes ranging between 600 to 700 for the different UCB references depending on filtering, whether nRBCs are included, and the combination of the references. The selected probes with annotation are provided in supplemental information (**Supplemental Tables S2 to 11**). Second, we used IDOL to select a L-DMR library which identified a L-DMR library consisting of 550 probes, of which 517 probes were present on both the 450K and 850K EPIC arrays, as the ideal number of probes for UCB deconvolution (**Supplemental Table S12)**.

The resulting L-DMR libraries from both methods were evaluated by calculating the R^2^ and root mean square error (RMSE) comparing estimates and FACS counts from each cell type in the test dataset. The results showing the pickCompProbes and IDOL optimized cell estimates precisions using the references as published and cleaned (using 70% cut off), both individually and combined, are summarized in **Figure 3** and presented in Supplemental **Tables S13 to 16**.

**Figure 3.**
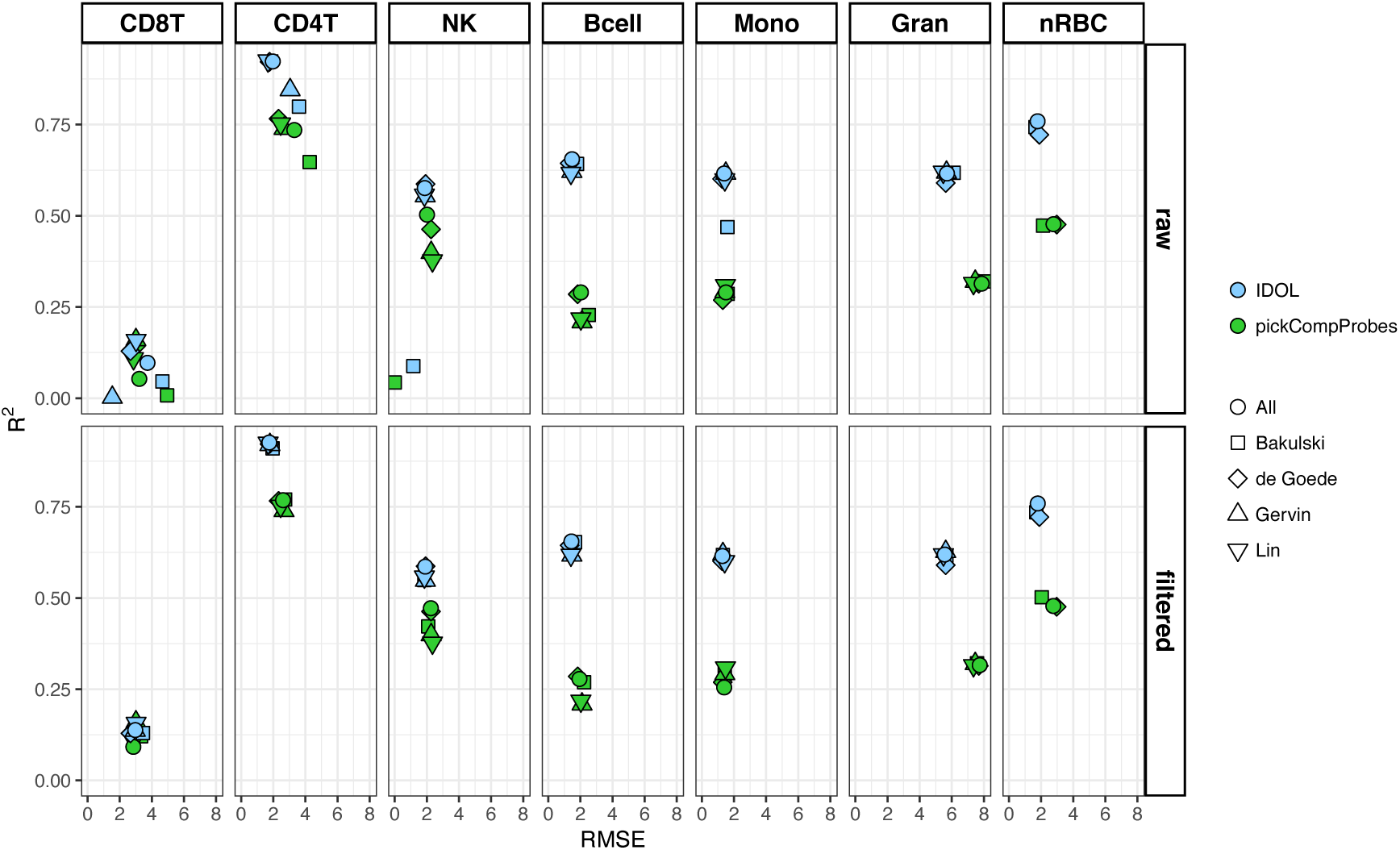
Evaluation of libraries. The selected libraries from pickCompProbes and IDOL were evaluated by calculating the R^2^ and RMSE comparing estimates and FACS counts from each cell type in the test dataset (n=22) using individual and combined UCB references. Mean R^2^ and RMSE are plotted on the y and x-axes, respectively.

The unfiltered *Bakulski reference* generated poorer cell type estimates (lower R^2^ and higher RMSE) compared to the other three reference datasets, though the filtered results were more similar across all individual references. Specifically, both pickCompProbes and IDOL performed poorly in estimating CD8T (pickCompProbes: R^2^=0.08, RSME=4.95 and IDOL: R^2^=0.46, RSME=4.66) and NK (pickCompProbes: R^2^=0.43, RSME=0 and IDOL: R^2^=0.88, RSME=1.16) proportions using the unfiltered *Bakulski reference*. Overall, the data cleaning and removal of contaminated samples significantly improved the accuracy of the deconvolution estimates. The R^2^ values were generally higher and RMSE values lower for the cleaned references, both individually and combined, across the two tested methods. Importantly, we obtained consistently more accurate estimates across all cell types when using the optimized IDOL L-DMR library compared to pickCompProbes (higher R^2^ and lower RMSE, **Figure 3**). This was observed across all individual reference datasets and in combinations. In fact, the R^2^ more than doubled for several cell types, particularly the estimated Bcell, Mono and Gran cell populations. Since the aggregated proportions of these cell types comprise approximately 70% of the leukocytes in cord blood, this is likely to have a substantial impact on the ability of the model to predict the “true” relative leukocyte composition. Similarly, the increased precision of the IDOL L-DMR to predict smaller proportions can reduce the confounding source of variability where subtle differences in cell types impact on the associations identified in EWASs. Combining all four references resulted in slightly and more consistently increased accuracy of the predictions across most of the cell types. Leaving out references one by one did not change the overall accuracy, although some cell type estimates were marginally improved. However, this was not consistent or attributable to one single reference dataset alone (**Supplemental Table S17**). In conclusion, these results clearly demonstrate the importance of using a clean combined reference and that the IDOL algorithm outperformed the pickCompProbes automatic selection of L-DMRs used for UCB deconvolution. We also investigated whether stricter cleaning (i.e. removal of samples instead of reclassifying to “correct” cell type) would impact our results, but found that this did not improve the accuracy of cell estimates (data not shown). Therefore, we based the downstream analyses upon the combined, cleaned UCB reference.

### Genomic location of the selected pickCompProbes and IDOL reference libraries

The probes selected by the two methods using combined, cleaned references are shown as heatmaps in **Figure 4A** and compared in **Figure 4B**. pickCompProbes did not discriminate as well as IDOL between hyper and hypomethylated probes in UCB (**Figure 4A**). Whereas the minfi L-DMR probes were biased towards hypermethylated, the IDOL L-DMR library consist of probes evenly distributed between hyper and hypomethylated. In addition, IDOL selected probes displaying intermediate DNAm, particularly for nRBCs. Only a small number of probes overlapped between the selections arising from applying pickCompProbes (to the raw and filtered references) and IDOL (n=45 and n=43, respectively). A comparison of the probes selected by the two methods with the genomic and functional context of the probes is provided in **Table 2**.

**Table 2.**
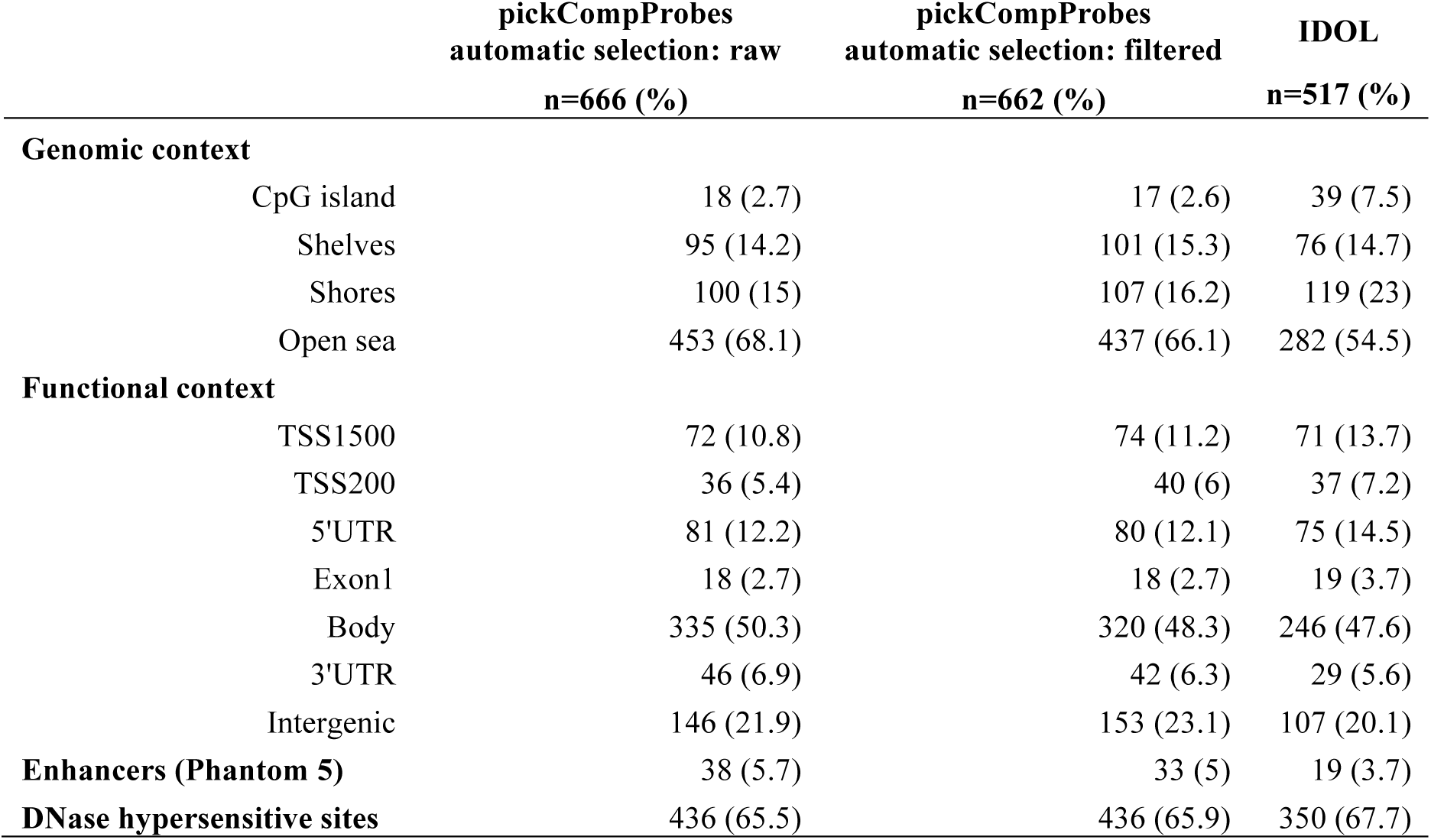
Genomic and functional context of IDOL and pickCompProbes (raw and filtered) libraries.

**Figure 4.**
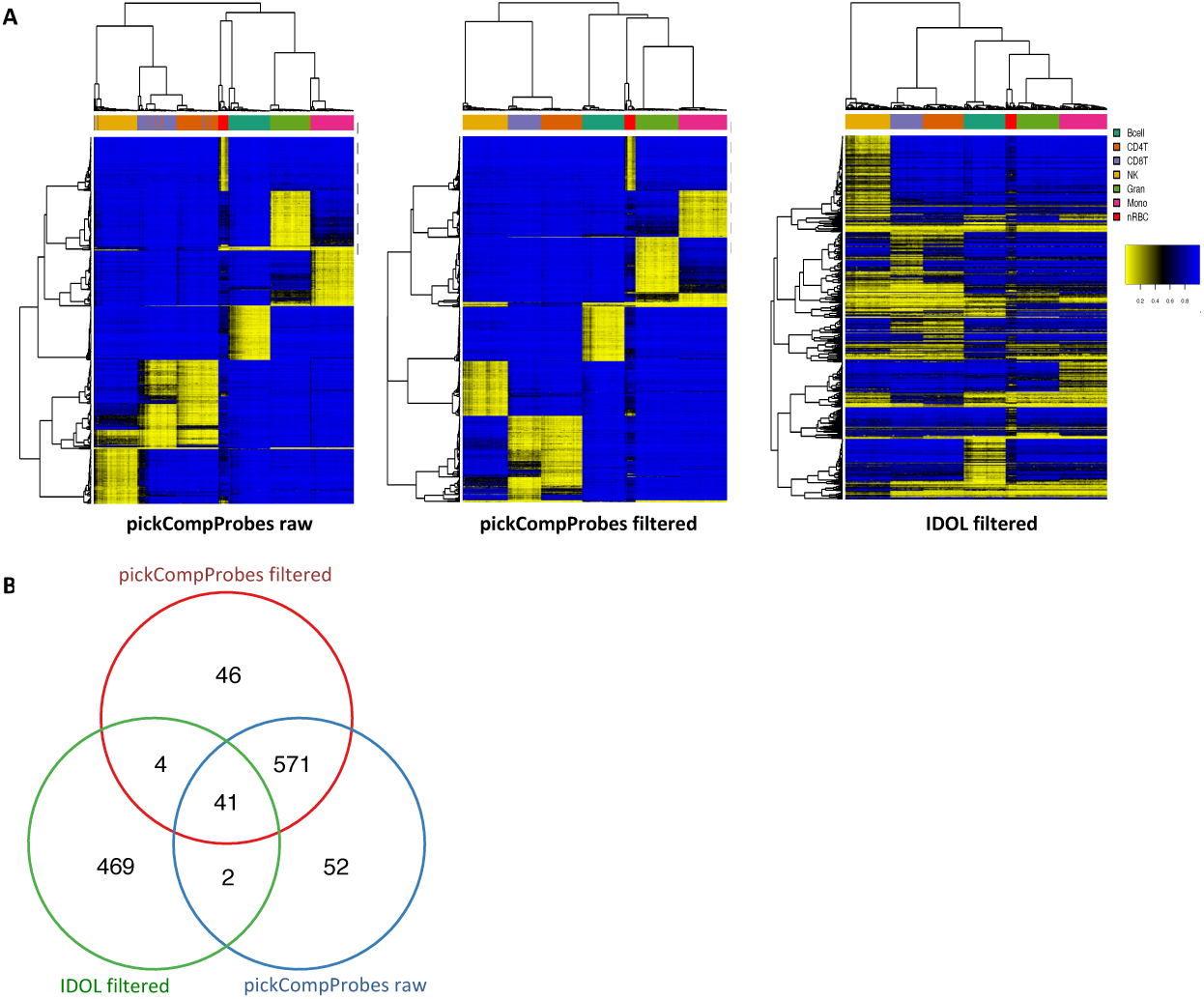
Comparison of L-DMR libraries selected using automatic selection in pickCompProbes and the IDOL algorithm for optimization. **A)** L-DMR libraries selected from combined UCB reference (raw n=666 and filtered n=662) using automatic selection in pickCompProbes and IDOL (n=517). **B)** Overlapping of probes from the three methods.

### Validation of deconvolution estimates in an independent birth cohort

Once we determined the L-DMR libraries for cell type estimation, we set out to estimate the accuracy of the deconvolution estimates in an independent cohort to assess the reproducibility of the reference datasets and selected libraries. To test this, we applied the CP/QP model using the pickCompProbes and IDOL L-DMR libraries to samples selected from the Generation R Study (n=191), from which cord blood 450K data and matched FACS cell counts were available[8, 23, 24]. Unfortunately, nRBC FACS cell counts were not available from the validation cohort, but approximately 90% of the total white blood cells were covered by the remaining 6 cell types in the references. Of note, the nRBC estimates were excluded from the predictions and the sum of the remaining six cell type estimates were rescaled to one so that the estimate proportions would be more similar to the FACS cell frequency data. Comparison of estimated cell type proportions with matched FACS counts revealed moderate R^2^ (coefficient of determination) values ranging from 15.19 to 78.86 % across cell type and method (**Figure 5**). Surprisingly, we obtained higher R^2^ for all cell types, except CD4T when using the L-DMR library generated with pickCompProbes compared to IDOL. However, the RMSE (root mean square error) was higher for the same cell types when using the pickCompProbes L-DMR, with a fold change in RMSE ranging from 0.93 to 2.33. Whereas the R^2^ is a relative measure of the fit, RMSE indicates the absolute fit of the model to the data (i.e. how close the FACS counts are to the deconvolution estimates predicted by the CP/QP model). RMSE is a measure of the accuracy of the model and the most important measure for the fit in this setting where the main purpose of the model is prediction. Hence, the consistently smaller RMSE using the IDOL L-DMR indicated a better fit and accuracy.

**Figure 5.**
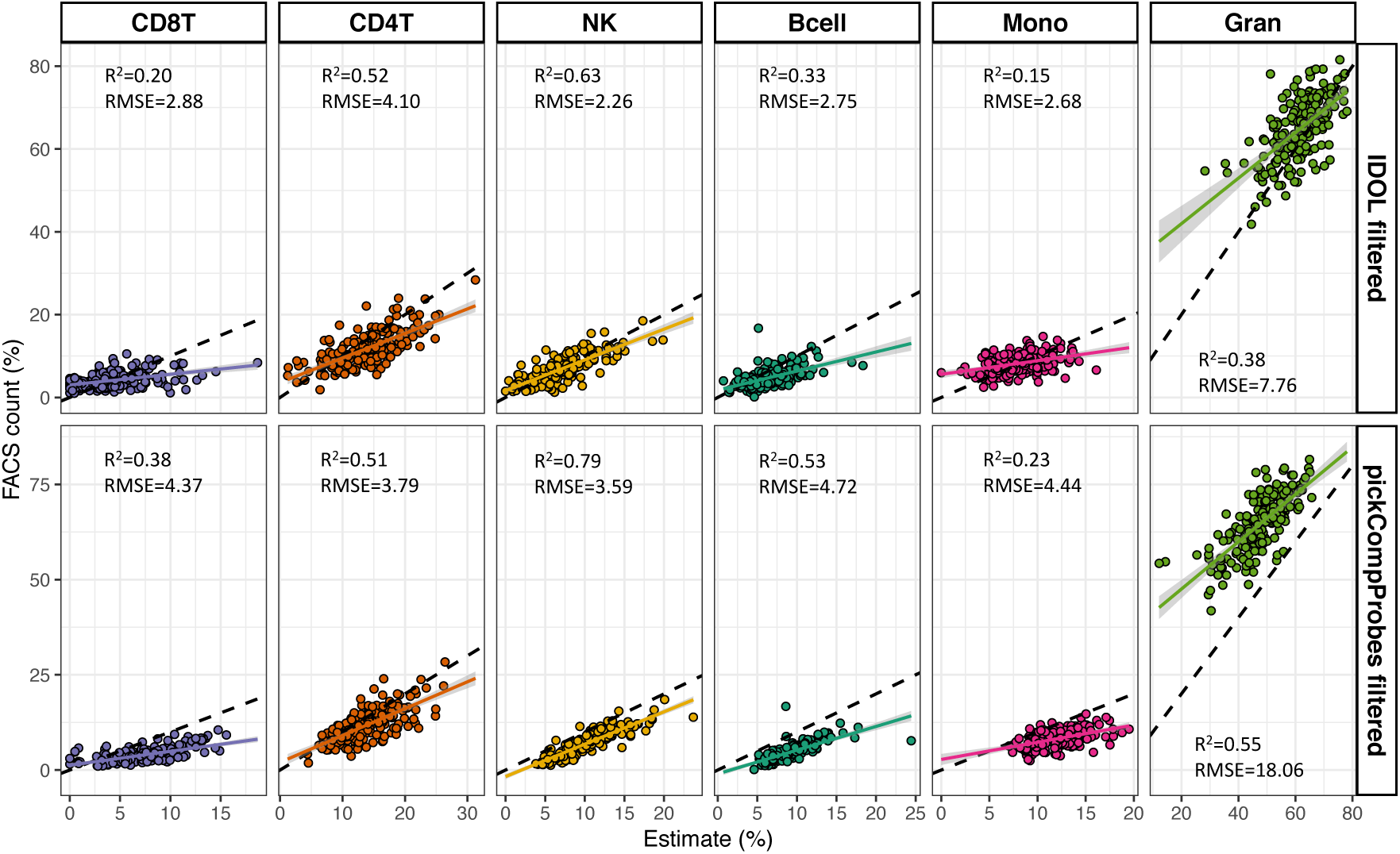
Comparison of estimated cell types and matched FACS cell counts. Scatter plots of deconvolution estimates using CP/QP programming and matched FACS cell counts in an individual birth cohort (Generation R, n=191) using cleaned IDOL and pickCompProbes libraries and the combined UCB reference. Smoothing lines represent the linear model. R^2^ and RMSE using the two methods are indicated for each cell type.

The higher R^2^ using pickCompProbes L-DMR showed how well the results adjusted to the line even if the line was shifted (higher correlation, but further away from the FACS counts). This was evident in the box plots of FACS cell counts and deconvolution estimates from the six cell types using the four methods (IDOL clean, IDOL clean strict, pickCompProbes using the filtered references and pickCompProbes using the raw references), which revealed a dramatic shift in the estimates using the pickCompProbes selection (**Figure 6A** and **Supplemental Figure S1**) compared to IDOL.

**Figure 6.**
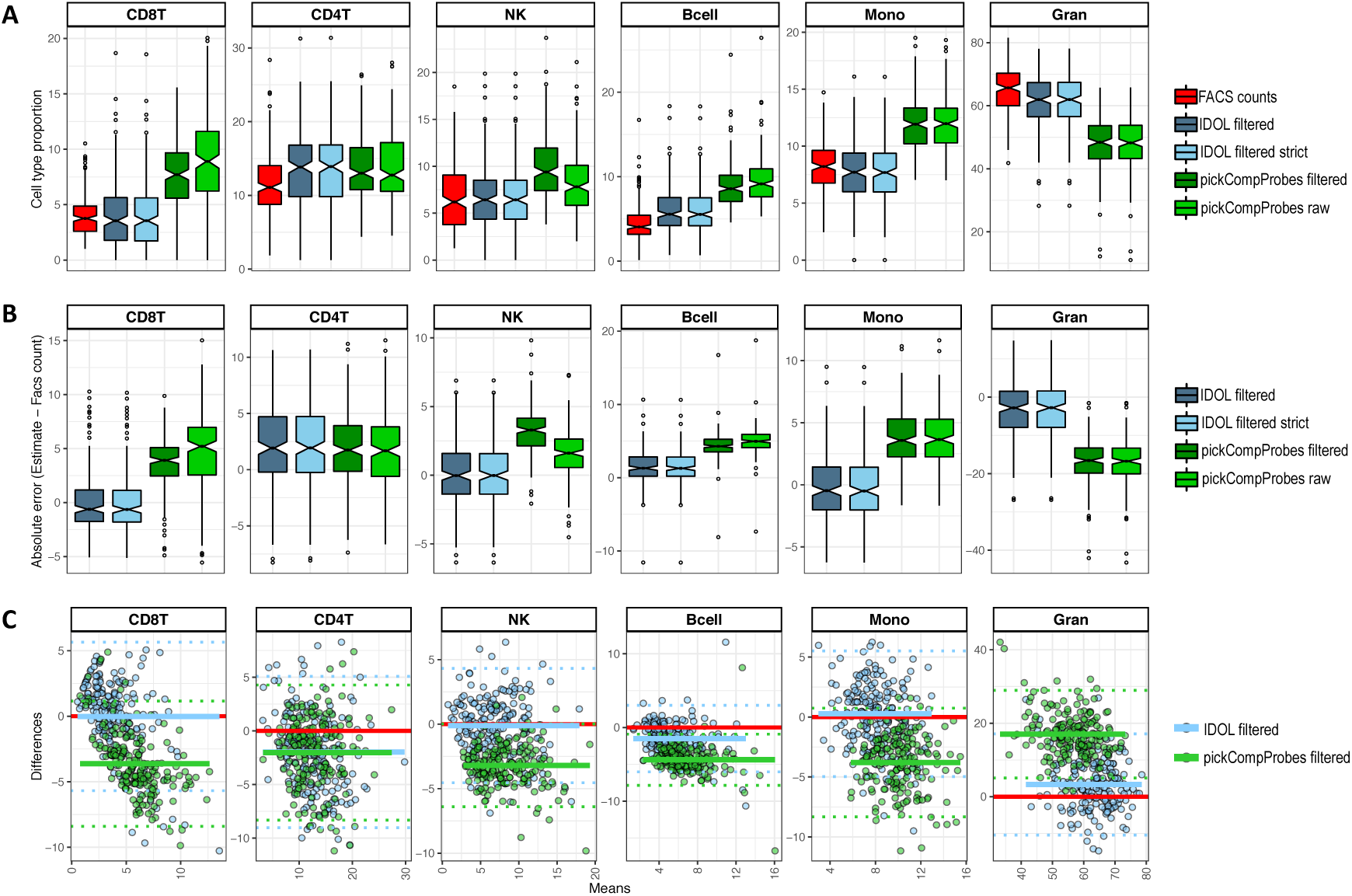
Measurements of accuracy and agreement between methods. **A)** Box plots of FACS cell counts (red) and estimates generated using IDOL (blue) and pickCompProbes (green) and a combined UCB reference (raw and filtered). **B)** Absolute errors (estimates minus FACS counts) by deconvolution method and the combined UCB reference (filtered and raw). **C)** Bland-Altman plots (differences versus means) showing the agreement between IDOL and pickCompProbes using a filtered combined UCB reference. The mean difference per method (blue and green) and zero difference (red) are indicated by horizontal lines.

Clearly, IDOL estimates approximated the actual proportions more precisely, independent of the cleaning process, while pickCompProbes estimates overestimated all cell types, except granulocytes. Further, subtracting the FACS cell counts from the deconvolution estimates (absolute errors) also shows that IDOL is more precise compared to pickCompProbes, which either over or underestimates the actual proportions (**Figure 6B** and **Supplemental Figure S2**). Finally, Bland-Altman plots (**Figure 6C**) comparing IDOL to pickCompProbes also show that IDOL results in more accurate UCB cell type predictions.

## Discussion

We conducted a systematic comparison and evaluation of four publicly available UCB reference datasets[17, 21, 22, 25] and provide recommendations for reference-based UCB deconvolution. We performed a descriptive characterization of the four datasets and determined the purity of the isolated cell fractions to generate clean references. To test the performance of raw and filtered references, both separately and in combinations, we applied two methods for selection of deconvolution libraries (i.e. automatic selection implemented in pickCompProbes and IDOL) and validated the deconvolution estimates in an independent cohort containing detailed, matched cell count information.

The UCB references were generated by four different laboratories and exhibit numerous technical and biological differences, which potentially impacts the downstream application of the data. Of these, purity and phenotyping of cells were assumed to have the largest impact on the resulting deconvolution estimates. Two frequently used methods for cell purification were used: MACS and FACS. Although relevant publications comparing MACS versus FACS related to cell enrichment, viability and functions of human blood cells were not available. FACS is known to often provide better purity[26]. Indeed, the purity estimates (if available) showed considerable variation between the UCB references.

We leveraged knowledge of blood cell types at a different developmental time point to implement a novel reference sample quality control process. We estimated the proportion of each of our UBC reference samples using an adult blood reference dataset derived from MACS sorted measures in men in mid-life. We looked for agreement between cell type identity in our UBC samples based on the adult measures. A filtering procedure (using an arbitrary cut-off estimating 70% of the purified cell type for inclusion) using a projection of adult cell types significantly improved the accuracy and precision of the deconvolution estimates. Although a 70% cut-off is moderate, we do not consider a stricter cut-off appropriate given the known physiological differences between cord and adult blood cell types. The cleaning resulted in removal of 29.6% of the samples from the *Bakulski reference*. Of note, almost all CD8 T cell fractions in this reference showed a large proportion of other cell types, mainly NK cells, when using a projection of adult CD8T cells. Further, 3 CD8T and CD4T samples from the same individual in the *Gervin reference* clustered to the opposite cell type. These samples were not processed together and we could not find evidence of sample swapping. Overall, the UCB CD4 and CD8 T cells displayed very similar DNAm patterns, whereas these cell types in adults differed more clearly [9, 19, 27]. The CD8T estimates using any of the UCB references were significantly less accurate compared to the other cell types, regardless of the library selection method. In general, UCB leukocytes are immunologically immature, as immunological memory still has to be established following responses to pathogens. The naive CD4 and CD8 T cells have not yet established their effector functions and this could explain why they are quite similar in their epigenetic profiles in UCB. Since T lymphocytes play a central role in adaptive immunity, CD4T and CD8T cells will eventually develop more cell type specific DNAm patterns as the cells mature[6, 7, 9].

Cell type specific DNAm analyses are challenging with respect to sample storage, processing, and cost, and all current reference datasets are limited by the number of available samples per cell type. Despite the range of biological and technical differences across the UCB references, combining references would yield a larger sample size, which we hypothesized would result in increased power and accuracy of the CP/QP model. One study effectively combined three reference datasets (de Goede, Bakulski and Gervin) for UCB deconvolution[28], though validating the specificity of a combined reference was not tested. Interestingly, although our analyses on combined data slightly improved accuracy and sensitivity of the deconvolution estimates, a clean reference reflecting cell type purity was much more important. Together, our findings demonstrate that reference datasets can be effectively combined across studies, enabling improved sample quality control and cell type estimation.

A critical first step in the CP/QP model is selection of suitable reference libraries (i.e. probes discriminating cell types). Two methods for library selection: the internal function pickCompProbes from minfi[16] and IDOL[18], were included for comparison in this study. The default selection of probes implemented in pickCompProbes performs differently for UCB (selecting sites agnostic of direction of methylation) and adult blood (selecting an even number of hyper and hypomethylated probes). Currently, this is the most frequently used method. In contrast, IDOL, which is an iterative algorithm, performs a dynamic scan and optimization of reference probes to accurately estimate cell types in a test sample set with known cell mixture values[18, 19]. It’s notable that the pickCompProbes and IDOL processes selected cell type distinguishing probe sets that were largely non-overlapping, underscoring the widespread genomic extent to which cell type specific DNAm patterns influence whole UBC measures. Following library selection via pickCompProbes or IDOL, the same CP/QP deconvolution methods for cell type estimation were performed.

We validated the combined references and library selection methods in an independent cohort (the Generation R Study), from which both matched, leukocyte-only FACS cell counts and UCB 450K data were available. Once we rescaled the CP/QP estimates to match the leukocyte proportions derived from the FACS cell counts, we observed results closer to the identity line using IDOL (lower RMSE and absolute errors). These analyses revealed that although we observed a better goodness of fit for pickCompProbes (higher R^2^ compared to IDOL), pickCompProbes estimates were biased with higher residual differences (and hence almost double RMSE compared to IDOL). This example shows the variance-bias trade-off between our IDOL L-DMR library compared to pickCompProbes [29]. pickCompProbes probe selection produces cell estimates with the lowest variance but shows higher bias. For investigators that seek relative cell proportion estimates within their population, these results suggest that pickCompProbes with the higher R^2^ is a preferred approach. In contrast, IDOL selects a L-DMR with the lowest bias, but as a trade-off increases the model variance (decreases the R^2^). For those investigators interested in predicting absolute cell proportions from DNA methylation data, as RMSE measures how accurately the model predicts the response, for this specific scenario with few outliers and a sample size >100, RMSE is the most important criterion for fit[30] and IDOL is the preferred estimation method.

Although we obtained good estimations using the individual cleaned references (**Supplemental Figures S1** and **S2**), unaccounted technical and genetic variability could increase residual noise during the estimations of specific cell subtypes. The use of average DNAm values across the four references within each cell type would reduce (though it will not eliminate) these sources of variability. Any difference in maternal age, gestational age, ethnicity, genetic polymorphisms and technical issues (e.g. during DNAm array processing) between the reference and validation cohorts may reduce the accuracy of the estimates. In addition, any difference in the fluorescent labeling of cells and performance of the conjugated antibodies between the two cell separation platforms (FACS and MACS) could contribute to the observed performance differences between individual references after cleaning. Specifically, in some references, the NK cells are not depleted for CD3, and thus might include a fraction of CD3+CD56+ T cells (also known as NKT cells) that could have affected the DNAm profile. Further, the NK cells in the validation cohort were depleted for CD3, but defined as being CD16 and/or CD56. Most of the CD16 cells are also CD56, but some are not, and could introduce more variation in the DNAm profile. As previously noted, it is not possible to unravel the impact these differences solely on the results presented here. However, as these subpopulations are very scarce, they will likely be absorbed within the next most similar cell population[19].

An additional limitation of the presented work is that only cell types in the reference datasets were estimated and validated. It is expected that cell types lacking reference DNAm signatures will be accounted for by the next most similar cell type represented in the reference dataset. In addition, different cell states and cell-cell interactions, which involves rapid epigenetic coordination, would not be picked up by a statistical model[7, 31]. Unmeasured cells are a potential weakness of any reference-based deconvolution method. Importantly, nRBC FACS counts were not available from this validation cohort. Approximately 90% of the total white blood cells however, were captured by the remaining six cell types in the references and validation datasets. nRBCs are a particular cell type of interest in the UBC community, given that their hypomethylation can be detected at half of the probes on the 450k array. A previous study tested one reference dataset (*Bakulski*) for validation with complete blood count cell measures in the Gen3G birth cohort. Among all pairwise estimated and measured cell types, they observed the highest correlation for nRBCs (R^2^=0.85)[32]. Given the reference sample overlap, we expect that our combined reference dataset would be applicable for nRBC estimation as well.

Based on our current findings, we recommend that studies with UBC DNAm measures on Illumina arrays who seek reference-based cell proportion estimates employ the following methods: 1) filtered and combined reference dataset available via Bioconductor as “*FlowSorted.CordBloodCombined.450k*”[33] and 2) *estimateCellCounts2()* function in the FlowSorted.Blood.EPIC R package[19], specifying all seven cell types, IDOL probe selection for deconvolution and noob preprocessing.

We look forward to expanded reference datasets available in the future for UCB and other tissues. Profiles generated in multiple laboratories with complementary methods, featuring cell types of expertise, provide a more comprehensive view of the tissue and time point. The development of multiple references also allows for reference sample quality control screening, with great improvement to estimates. Here, we demonstrate the utility of developing a cross-lab reference dataset to harness the power of multiple studies. While UCB is a well-characterized tissue, advances in single-cell sequencing technology will enable future characterization and identification of rare and intermediate cell-states.

## Conclusions

In conclusion, these results clearly demonstrate the importance of using a filtered, combined reference for estimating cell proportions from DNAm data. The IDOL library selection significantly outperformed the pickCompProbes automatic selection of probes used for UCB deconvolution. These results have important implications for future epigenome-wide studies of DNA methylation in cord blood, offering a method that has the potential to reduce confounding due to cellular heterogeneity. Further, we provide guidelines for reference-based cord blood deconvolution.

## Methods

### Description of datasets

Four publicly available UCB reference datasets were included in this study, here named by first author of the respective papers and listed by time of publication: *de Goede reference*[25], *Bakulski reference*[17], *Gervin reference*[21] and *Lin reference*[22]. The first three datasets consist of 450K DNAm data and the Lin reference contains EPIC 850K DNAm data. Validation and benchmarking of the estimated cell type proportions to matched cell counts was performed in independent UCB samples selected from the Generation R study[24, 34].

#### de Goede reference

The de Goede reference is obtained from UCB samples (n=7, five female and two male) collected at the BC Women’s Hospital. From each sample, seven cell types (Gran, Mono, Bcell, CD4T, CD8T, NK and nRBCs) were separated using FACS. Of note, one participant’s sample did not result in sufficient CD8T cell DNA and so was not included. In addition, the DNAm profiles from whole cord blood prior to separation were included. 450K DNAm data was retrieved from the GEO data repository (GEO accession GSE68456). For full details, refer to de Goede et al.[25].

#### Bakulski reference

The Bakulski reference was obtained from UCB samples (n=15, 8 male and 7 female) collected from full-term deliveries at John Hopkins Hospital. From each sample, seven cell types (Gran, Mono, Bcell, CD4T, CD8T, NK and nRBCs) were separated using MACS and double density centrifugation. In addition, the DNAm profiles from whole cord blood prior to separation were included. The race/ethnicity of the participants are unknown. The resulting dataset consisted of a variable number of cell type fractions included in the final data. 450K DNAm data was retrieved from the R package *FlowSorted.CordBlood.450K* in Bioconductor. For full details, refer to Bakulski et al.[17].

#### Gervin reference

The Gervin reference was obtained from UCB samples (n=11, 5 male and 6 female) from full-term births at the Alternative Birth Care unit at Oslo University Hospital in Norway. The Gervin reference consisted of all native Norwegians. All samples were successfully fractionated into six cell types (Gran, Mono, Bcell, CD4T, CD8T and NK) using FACS. Notably, nRBCs were not included in this reference dataset. 450K DNAm data was retrieved from the R package *FlowSorted.CordBloodNorway.450K* in Bioconductor. For full details, refer to Gervin et al.[21].

#### Lin reference

The Lin reference was obtained from UCB samples (n=14, 10 male and 4 female) from full-term births. In addition, this reference also contains cord blood tissue, which has not been included for comparison in the present study. The ethnic distribution of subjects included 5 Chinese, 4 Malay and 5 Indian. All UCB samples were fractionated into six cell types (Gran, Mono, Bcell, CD8T, CD4T and NK) using MACS. In addition, UCB buffy coat samples were also assayed for DNAm prior to fractionation. Notably, nRBCs were not isolated in this reference dataset. EPIC 850K DNAm data from UCB was retrieved from the R package *FlowSorted.CordTissueAndBlood.EPIC*. For full details, refer to Lin et al.[22].

#### Jones dataset (test dataset)

For IDOL optimization and for comparison versus the pickCompProbes procedure, we used a data set (GEO GSE127824, n=24) collected at the British Columbia Women’s hospital consisting of UCB DNAm signatures and matched FACS counts for the seven cell types with available references (Gran, Mono, Bcell, CD4T, CD8T, NK and nRBCs). One sample showed extreme FACS values for NK (14%) and a second sample showed low granulocytes (23%) and high CD4T (51%). As these extreme values could influence the R^2^ results in our comparisons, we restricted the samples to n=22.

#### Generation R dataset (validation cohort)

Immunophenotyping of white blood cell subsets using FACS and whole UCB DNAm measures in the samples (n=191) selected from the Generation R study are described elsewhere[35].

#### Adult reference

The adult blood reference dataset correspond to the recently published FlowSorted.Blood.EPIC library[19]. Briefly, six MACS-isolated and FACS-verified pure cell subtypes (neutrophils (Neu), Mono, Bcell, CD4T, CD8T, and NK) were purchased from commercial vendors. Cells were isolated from 31 males and 6 females, all anonymous healthy donors. The donors had a mean age of 32.6 years (range 19–59 years) and an average weight of 86 kg (range 65–118 kg) and were negative for HIV, HBV, and HBC. Women were not pregnant at the time of sample collection, and samples were collected from donors with no history of heart, lung, or kidney disease, asthma, blood disorders, autoimmune disorders, cancer, or diabetes. DNAm was measured using Illumina HumanMethylationEPIC array. For our analyses, we used the legacy IDOL library optimized for Illumina HumanMethylation450k using artificial mixtures as described in the original paper[19].

### Data processing and filtering

#### Data processing

All analyses were carried out using the R programming language (http://www.r-project.org/). DNAm data from all datasets were preprocessed in minfi[16] using *preprocessNoob* for background and dye-bias normalization. We performed a general quality control to ascertain that none of the samples showed signs of bisulfite conversion problems or hybridization technical defects. None of the samples showed more than 5% detection p values over the background >10E-07. We did not have access to the number of beads information for most of these samples, thus no additional filtering using that criterion was performed. Although we examined strict filtering excluding X and Y-chromosomes and potential cross-reactive probes, we decided to use the complete information from the raw datasets without any filtering. Given potential technical batch issues when combining the two different array platforms (450K and EPIC with different intensity ranges), we first used the combined 450K datasets for IDOL optimization. This resulted in a dataset consisting of 207 samples for the Bakulski, de Goede and Gervin references (see **Table 1**). After data cleaning and IDOL optimization we included the Lin dataset (n= 83). We observed 452 567 overlapping probes shared across all four reference datasets and array technology platforms.

#### Reference data sample filtering

Each reference dataset possesses technical and biological differences, each with potential strengths and weaknesses. Differences in purity estimates and phenotyping of cell types will likely influence the deconvolution estimates from the datasets individually, and particularly in combination. To test this, we reasoned that a strict cleaning of the datasets prior to deconvolution could improve the accuracy and precision of the deconvolution estimates. Specifically, we projected adult cell types (from the FlowSorted.Blood.EPIC R package[19]) onto the sorted UCB reference samples. We visualized the relative proportions of each adult cell type that were predicted in the cord sample using stacked bar charts. We compared the observed adult cell type estimates the expected cord cell type identity with an arbitrary inclusion cut-off of 70% of the expected purified cell type. Consequently, any sample with <70% of the equivalent adult cell type was considered a cell mixture and removed from the reference. Samples with >70% of a different cell type were reclassified to the “correct” cell type. This quality control step was applied to UCB cell types with parallel adult cell types available (Gran, Mono, Bcell, CD8T, CD4T and NK). Due to adult and infant physiologic differences, this step was not performed for nRBCs.

### Descriptive comparison of reference datasets

To compare the UCB references, we used multidimensional reduction and unsupervised hierarchical clustering. For dimensionality reduction, we performed PCA using the *prcomp* function in stats R package. In each reference dataset, we computed the variance explained by the principal components. We plotted principal components one and two, colored by cell type label, and visually inspected for overlap of cell types.

### Selection of reference libraries and deconvolution

We used two approaches for selection of reference libraries prior to applying the Houseman CP/QP method implemented in minfi for deconvoluting Gran, Mono, Bcell, CD4, CD8, NK and nRBCs.

#### Automatic probe selection (pickCompProbes)

We used the *estimateCellCount*s*2* function in the FlowSorted.Blood.EPIC R package[19] to automatically select library probes from each reference data separately and in combinations, by specifying the default cord blood options. The *estimateCellCount*s*2* function is a modification of the commonly used *estimateCellCounts* function in minfi[16]. Notably, *estimateCellCount*s*2* provides a flexible input of any RGChannelSet or raw MethylSet and selection of a customized set of probes obtained from IDOL optimization in addition to the pickCompProbes included in the original function. According to previous literature, we used the automatic selection (option “any”), choosing the top 100 most differentially methylated probes per cell type based on F-test statistics. Depending on the library, the number of probes could be 700, or fewer if some of the selected probes overlapped.

#### IDOL algorithm

A complete explanation of the IDOL algorithm is described elsewhere, please refer to Koestler et al[18] and Salas et al[19]. In brief, the IDOL algorithm performs a dynamic search in a candidate set of cell type specific DNAm markers for a library that is optimized to accurately estimate cell types. In the IDOL algorithm, a series of two-sample t-tests are first fit to the matrix of CpGs and used to compare the mean methylation beta-values between each of the *K* cell types against the mean DNAm beta-values computed across the remaining *K* - 1 cell types. Putative L-DMRs are identified by first rank ordering CpGs by their *t*-statistics, then taking the top *M* (150 CpGs in our process) L-DMRs with the smallest and largest *t*-statistics for each of the *K* comparisons. Using this list, IDOL performs an iterative process trying to find the optimal probes (highest R^2^ and lowest RMSE) for a defined size of candidates. The procedure collects the optimal candidates for several predefined L-DMR library sizes. To define optimal library the algorithm compares the cell estimates obtained using CP/QP in each iteration versus the true estimates in the mixture (e.g. the cell proportions derived from the FACS cell counts).

IDOL optimization was performed using the Jones dataset (n=22). The Jones dataset consists of UBC samples in which the “true values” for the seven cell subtypes interrogated in the reference libraries is known (FACS information). The samples in this dataset were divided half into training (n=11) and half into testing (n=11) groups for the IDOL L-DMR libraries. Specifically, to calibrate the selection of optimal L-DMR libraries, we applied IDOL to the training set to identify several sets of optimized reference libraries for UCB deconvolution. Libraries ranging from 300 to 600 probes have previously demonstrated to generate accurate and reliable deconvolution estimates[19]. Hence, we chose a preselected library size between 150 to 700 probes with increments of 50 probes per each optimization cycle. During the implementation of the IDOL a total of 500 iterations were run for library optimization. The IDOL reference libraries were evaluated by calculating the R^2^ and RMSE comparing the proportion estimates using CP/QP and the proportions derived from the FACS counts for each cell type in the 11 samples used for training versus the 11 samples used for testing each library. Among the 12 sets of libraries the optimal library showed the highest R^2^ and the lowest RMSE on both the testing and training sets when comparing the cell estimates versus the proportions derived from the FACS counts.

#### Comparison of selected libraries

We identified four libraries (pickCompProbes raw, pickCompProbes filtered, IDOL raw, and IDOL filtered). We used heatmaps with hierarchical clustering to visualize sorted cell type methylation levels in each of the libraries selected. We used an Euler diagram to describe the relationships between the probes selected for the library methods. The selected libraries were used for deconvoluting Gran, Mono, Bcell, CD4T, CD8T, NK and nRBCs by applying the Houseman CP/QP method to obtain cell type estimates in the test samples (n=22). We used the combined reference dataset as well as the four reference datasets individually. The cell proportion estimates resulting from these libraries and reference datasets were compared to FACS counts for each cell type and evaluated by calculating the R^2^ and RMSE.

#### Leave-one-out analysis

We compared the cell estimates obtained using the IDOL procedure using the combination of the four references versus those obtained excluding one of the references while leaving the other three.

## Validation of the deconvolution estimates

The accuracy of the deconvolution methods was validated by comparing pickCompProbes and IDOL cell type estimates against matched cell counts in cord blood samples selected from the Generation R study (n=191). Note, nRBCs were not available in the validation dataset and testing was performed exclusively on CD8T, CD4T, NK, Bcell, Mono, and Gran cell types. Correlation of the cell type estimates and matched cell counts was calculated using the Pearson correlation test. We further calculated the absolute errors (cell type estimates minus FACS counts) by library selection method and by the combined UCB reference (filtered and raw). With the filtered, combined UCB reference dataset, we used Bland-Altman plots to assess the difference versus the mean cell type estimate per library selection method (pickCompProbes and IDOL).

## List of abbreviations

UCB: umbilical cord blood
DNAm: DNA methylation
IDOL: Identifying Optimal Libraries
EWAS: epigenome-wide association studies
nRBCs: nucleated red blood cells
CP/QP: constrained projection/quadratic programming
L-DMR: Leukocyte differentially methylated regions
PCA: principal component analysis
MACS: Magnetic-Activated Cell Sorting
FACS: Fluorescence-Activated Cell Sorting
RMSE: root mean square error
CD8T: CD8^+^ cytotoxic T-lymphocytes
CD4T: CD4^+^ helper T-lymphocytes
NK: Natural Killer cells
Bcell: B-lymphocytes
Mono: Monocytes
Gran: Granulocytes

## Declarations

### Ethics approval and consent to participate

The Generation R Study was approved by the medical ethics committee of Erasmus MC, University Medical Center Rotterdam. Informed consent was obtained for all participants.

### Consent for publication

Not applicable

### Availability of data and materials

The Jones dataset (test dataset) included the current study is available in GEO GSE127824 (https://www.ncbi.nlm.nih.gov/geo/query/acc.cgi?acc=GSE127824). FlowSorted.CordBloodCombined.450k is available in Bioconductor (https://doi.org/doi:10.18129/B9.bioc.FlowSorted.CordBloodCombined.450k) and the original source code is available through https://github.com/immunomethylomics/FlowSorted.CordBloodCombined.450k (under license GPL-3.0). For reproducibility the source code has also been deposited on Zenodo (doi: https://doi.org/10.5281/zenodo.2584162 for the package and doi: https://doi.org/10.5281/zenodo.2584381 for the scripts used in the analyses).

### Authors’ contributions

KG collected the Gervin reference data, conceived the study, performed data analysis and drafted the manuscript. LAS conceived the study, performed data analysis, and drafted the manuscript. KMB collected the Bakulski reference data, advised on data analysis and helped edit the manuscript. MCvZ generated FACS data for the GenR cohort. DCK advised on data analysis and helped edit the manuscript. JKW advised on data analysis and helped edit the manuscript. LD generated FACS data for the GenR cohort. HAM generated FACS data for the GenR cohort. KTK advised on data analysis and helped edit the manuscript. MSK conceived the de Goede reference data and helped edit the manuscript. RL helped conceive the study and edit the manuscript. BCC advised on data analysis and helped edit the manuscript. JF conceive the study, performed validation analysis in the Generation R Study, advised on study design and helped edit the manuscript. MJJ conceived the study, performed data analysis and drafted the manuscript. All authors read and approved the final manuscript.

## Acknowledgements

The Generation R Study is conducted by the Erasmus Medical Center in close collaboration with the School of Law and Faculty of Social Sciences of the Erasmus University Rotterdam, the Municipal Health Service Rotterdam area, Rotterdam, the Rotterdam Homecare Foundation, Rotterdam and the Stichting Trombosedienst & Artsenlaboratorium Rijnmond (STAR-MDC), Rotterdam. We gratefully acknowledge the contribution of children and parents, general practitioners, hospitals, midwives and pharmacies in Rotterdam. The study protocol was approved by the Medical Ethical Committee of the Erasmus Medical Centre, Rotterdam. Written informed consent was obtained for all participants. The generation and management of the Illumina 450K methylation array data (EWAS data) for the Generation R Study was executed by the Human Genotyping Facility of the Genetic Laboratory of the Department of Internal Medicine, Erasmus MC, the Netherlands. We thank Mr. Michael Verbiest, Ms. Mila Jhamai, Ms. Sarah Higgins, Mr. Marijn Verkerk and Dr. Lisette Stolk for their help in creating the EWAS database.

## Supplemental information

**Figure S1.**
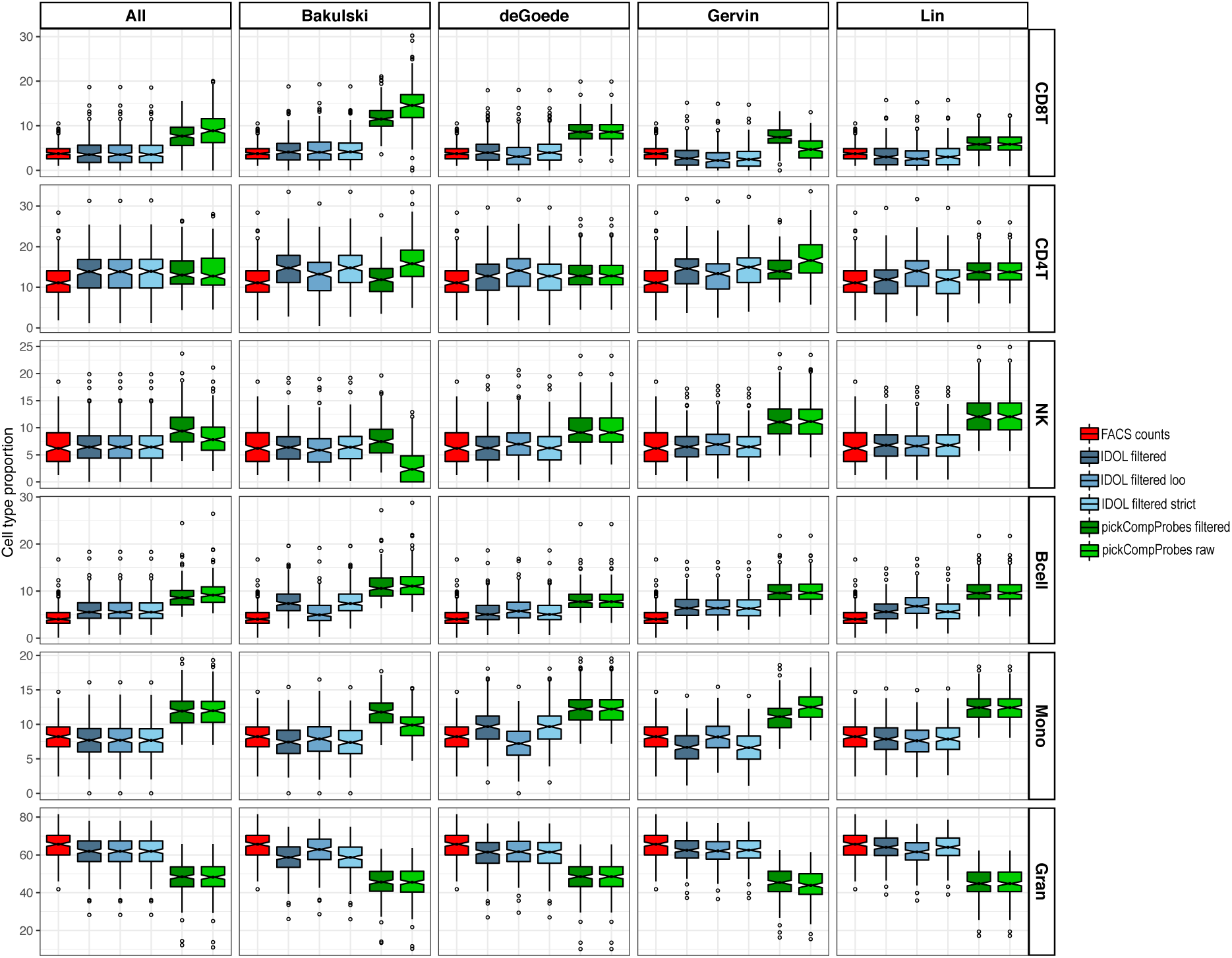
Box plots of estimates and FACS counts. Cell type proportions generated by FACS (true values) and CP/QP programming (estimates) using IDOL and pickCompProbes L-DMRs for each UCB reference individually and combined.

**Figure S2.**
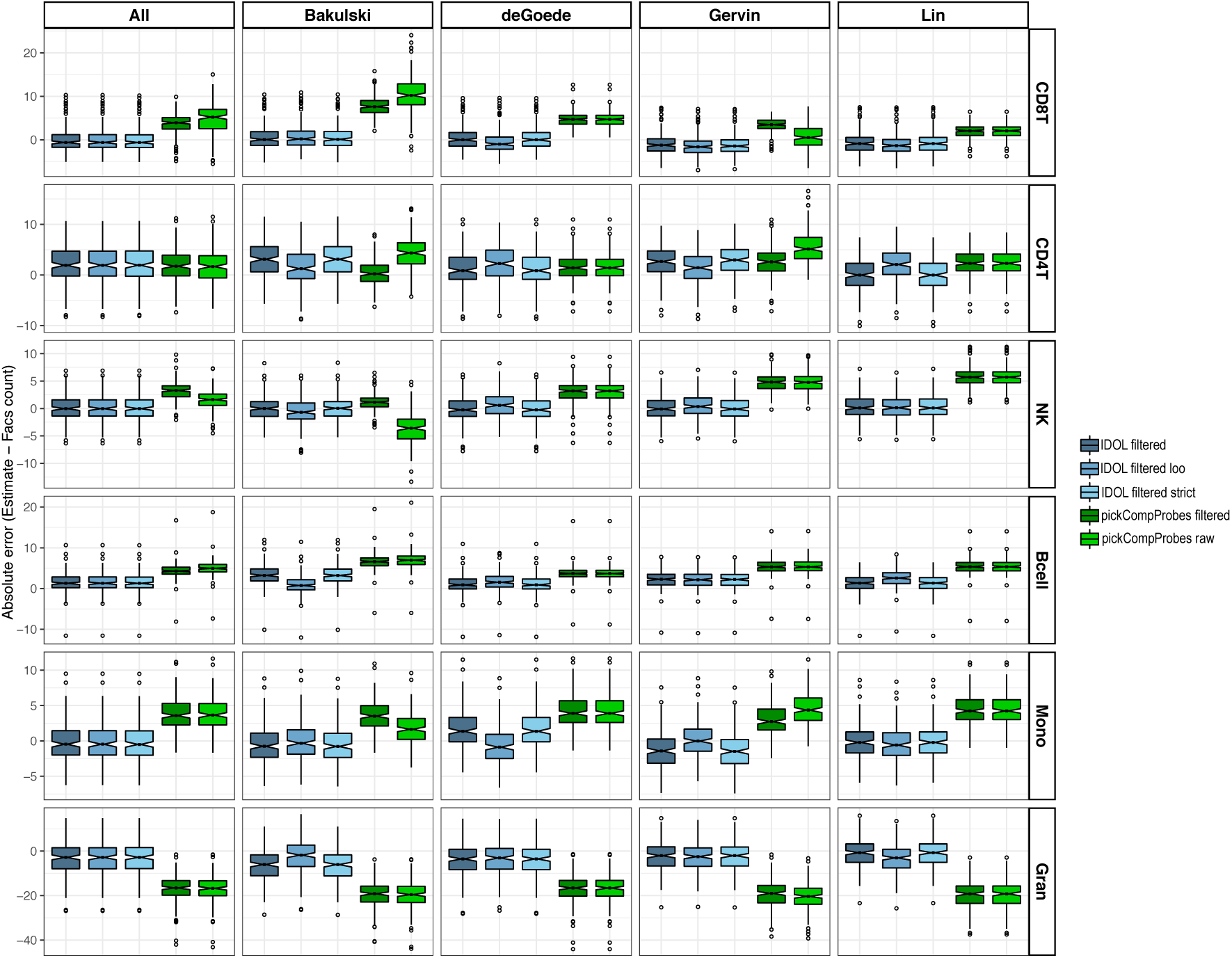
Box plots of absolute errors. Absolute errors (estimates minus FACS counts) per method (IDOL and pickCompProbes) for each UCB reference individually and combined.

